# Individual differences of cortical and subcortical emotion-informed functional gradients

**DOI:** 10.64898/2026.02.09.704784

**Authors:** Chun Hei Michael Chan, Laura Vilaclara, Patrik Vuilleumier, Dimitri Van De Ville, Elenor Morgenroth

## Abstract

The complex interplay between brain regions that support emotional experience and their link to individual differences is a topic of active research. Additionally, there has been growing interest in using functional gradients to investigate human cortical organization during both rest and film fMRI. Among these, several studies demonstrated improved brain fingerprinting performance, reflecting greater neural identification capability of film fMRI against rest fMRI despite higher subject synchronization during film-watching than in rest. Comparably, in this work we study the relation between individual differences, in particular, state anxiety and openness scores, and brain activity during the processing of various emotional scenes in films, through functional gradients. Next to including subcortical areas, we also propose a new approach of computing functional gradients based on a subset of frames selected using emotional annotation data of films, resulting in emotion-informed functional gradients. Then we evaluate the variance in emotion-informed gradients across subjects and employ these same gradients in the prediction of individual differences. For emotion-informed functional gradients, the highest predictability of state anxiety was found for scenes of negative valence and medium-high arousal, corresponding to the typical location of anxiety within the valence-arousal-power emotional space. Additionally, predictability of state anxiety was negatively correlated to inter-subject variability. In contrast, predictability of openness was found to be highest during scenes with low arousal and positively correlated to inter-subject variability. In essence, our results first show that macroscale brain organization is affected by emotional experience, and that frame selection based on the latter can be useful to remove non-subject-specific variability while extracting subject-specific information related to the emotion experience. It also demonstrates that frame selection increases inter-subject variability allowing the extraction of more subject-specific information. Thus, expanding on the idea of brain fingerprint in film fMRI, we argue that emotional experiences enhance disentanglement of various domain of individual differences. Moreover, depending on the individual difference of interest, fMRI acquired during more or less constrained paradigms would be more suitable to reveal different properties of brain function.

## 1. Introduction

During the past three decades, functional magnetic resonance imaging (fMRI) has become a tool of choice in neuroscience to indirectly probe human brain activity. The analysis of fMRI blood-oxygenation-level-dependent (BOLD) signals has long been focused on univariate approaches that contrast activation over different task blocks to ascribe specific brain areas to distinct functions.

Another viewpoint emerged along with the advent of resting-state fMRI, leveraging the concept of functional connectivity (FC); i.e., by computing the temporal correlations between brain regions, we can understand how regions interact and constitute distributed functional networks (Biswal et al., 1995; Fox et al., 2005). This approach has revealed the intrinsic organization of brain activation in the absence of task and made it possible to associate changes in this organization with individual differences or diseases. Therefore, resting-state fMRI and the advanced analysis techniques that came with it offer a major breakthrough in neuroscience (Finn, 2021). However, measuring brain activation in the absence of task or stimulation has also its limitations, and brain activity patterns vary with emotional state, including during resting state (Eryilmaz et al., 2011; Gaviria et al., 2021b).

The use of film in fMRI (Morgenroth et al., 2023b; Hasson et al., 2008) has recently gained great popularity, and many of the analysis techniques used for it have their origins in resting-state fMRI. Film fMRI has been shown to have numerous advantages over resting-state fMRI, such as increased participant compliance, higher test-retest reliability, but also improved decoding of individual differences (Finn and Bandettini, 2021; Vanderwal et al., 2017, 2019; Samara et al., 2022). For example, when using FC to predict behaviour and subject identity, film watching was shown to be superior to task conditions (Finn and Bandettini, 2021). Films contain highly social and emotional content, thus they can evoke brain activation related to richer processes that may be highly individual. This has been proposed as a reason why film outperforms rest for predicting individual differences (Finn and Bandettini, 2021).

In addition to the study of individual differences and subject identification, film fMRI has been used in affective neuroscience to study brain activation related to emotion experience. Numerous studies have applied machine learning techniques to predict affective content in films from fMRI data (Kragel et al., 2018; Kragel and LaBar, 2015; Horikawa et al., 2020; Saarimäki et al., 2016; Kim et al., 2019). These studies often rely on mass-univariate models or FC for prediction (Mohammadi et al., 2023; Gaviria et al., 2021a,b). This work shows that film fMRI not only contains subject-specific information, but also rich stimulus-driven information that goes beyond the sensory features of audio-visual stimulation.

A major feature of film fMRI compared to rest is that brain activation between subjects gets synchronized (Hasson et al., 2004). Inter-subject synchronization is strongest in brain regions associated with audio-visual processing, yet it occurs across the whole brain. Importantly, when investigating inter-subject synchronization dynamically, it has been shown that it is especially strong during highly social and emotional events (Li et al., 2021; Gruskin et al., 2020; Bolton et al., 2018). The surprising relationship between increased inter-subject synchronization during film fMRI and improved predictability of individual differences compared to rest fMRI has not been directly investigated. Here we consider the effect of emotion, modulating subject synchronization and predictability of individual differences in the fMRI signal during film viewing.

We employ functional gradients, which are a lower-dimensional connectivity representation of the macrostructure of brain organization. Functional gradients, first introduced in 2016 by (Margulies et al., 2016) for resting state, have already been used in film fMRI (Samara et al., 2022; Huntenburg et al., 2018). They describe a low-dimensional approximation of FC matrices in terms of connectivity profiles; i.e., two points (representing regions) close to each other in the gradient space have similar connectivity profiles. Similar to measures of FC, it has been demonstrated that also when using functional gradients (Samara et al., 2022), film fMRI provides a better subject behaviour predictability over resting-state fMRI.

In this study, our objective is to examine how functional gradients reorganize during the processing of emotional scenes in films by investigating inter-subject variability and predictability of individual differences. Specifically, we first replicate gradients’ layout for films and rest as previously shown Samara et al. (2022). We observe inter-subject variability and predictability of individual differences in film and rest gradients. Then, we examine the organization of emotion-informed gradients constructed by leveraging timepoints that are situated within specific areas of the Valence-Arousal-Power (VAP) emotion space Mehrabian and Russell (1974), a three-dimensional model that characterizes emotional experiences along the dimensions of valence (ranging from pleasant to unpleasant), arousal (ranging from calm to excited), and power (reflecting the sense of control or dominance). We additionally verify their test-retest reliability across films, showing that emotion-informed gradients yield similar results across timepoints belonging to the same area of the emotion space, despite subjects being effectively shown different scenes. We then use the emotion-informed gradients to predict individual differences. In particular, we locate the most predictive areas of the VAP emotion space, i.e which type of emotional scene associated gradient is most predictive. Finally we relate the resulting predictability to inter-subject variability using correlation across areas of the VAP emotion space. Through this last analysis, we provide further insight into the interplay between subject synchronization and predictability of individual differences.

We hypothesize that film fMRI will provide greater predictive power of individual differences than rest fMRI, and we seek to determine how this advantage is modulated by affective processes. We further expect that the inter-subject variability of functional connectivity (FC) gradients will be higher during rest than during film viewing, consistent with the stronger within-individual alignment of large-scale brain connectivity under naturalistic stimulation. Finally, we will tie these findings together by examining the relationship between predictive performance and inter-subject variability in FC gradients during film fMRI.

## 2. Methods

### 2.1. Data

We used the Emo-FiLM dataset, which includes data from a behavioural and an fMRI study of film watching. Data acquisition, preprocessing, and quality is described in detail in (Morgenroth et al., 2024). From this dataset, we employ fMRI recordings from 30 healthy participants (18 female, average age=25.83, std=3.60) during a ten-minute resting-state scan and while watching 14 short films.

#### MRI Acquisition and Preprocessing

All fMRI images were acquired on a 3T Siemens Magnetom TIM Trio scanner at the Brain & Behavior Laboratory (BBL) of University of Geneva, with the same simultaneous multi-slice gradient-echo planar imaging sequence provided by the Centre for Magnetic Resonance Research (CMRR, Minnesota) [37], [38] (TR = 1.3s, TE = 30ms, flip angle = 64°, MB acceleration factor = 3, interleaved MB mode, 54 slices, FoV read = 210mm, voxel size = 2.5 x 2.5 x 2.5 mm^3^, PE = AP, bandwidth = 2290 Hz/Px, Echo Spacing = 0.57ms, EPI factor = 84, Pulse duration = 4300us, fat saturation). Resting-state runs lasted 10 minutes, totalling 460 volumes. Film runs differed in duration, as film duration ranged from 6:42 to 17:08. Film runs included a 90s rest period before and after each film. Preprocessing was performed using FEAT (FMRI Expert Analysis Tool) Version 6.00, part of FSL and included registration, motion correction, spatial smoothing (Gaussian Kernel of FWHM 6mm) and temporal filtering (Gaussian-weighted least-squares straight line fitting, with sigma = 50 s) to prevent spurious fluctuations from our dynamic window approach (Leonardi and Van De Ville, 2015). Furthermore, six motion regressors and average time courses of WM and CSF were regressed from the data.

In addition to the preprocessing performed in the Emo-FiLM dataset, we implemented scrubbing, removing time points with framewise displacement exceeding 0.5 mm. Furthermore, we extracted region-averaged timecourses for the Schaefer 400 functional parcellation and 14 subcortical areas from the Harvard Oxford Subcortical Atlas, resulting in timecourses for 414 brain regions for each run. For network-level analysis, we split these regions into a total of eight functional networks comprising of the seven networks from (Thomas Yeo et al., 2011) and adding the subcortical regions. Specifically for films, rest periods that were at the beginning and end of each film run were removed under consideration of a lag to account for the hemodynamic response function. We shifted the timecourses by 4 TRs (≈ 6 seconds) to only include TRs with activation corresponding to film watching in our analysis of film runs.

#### Emotion Experience Timecourses

We used consensus annotations of experienced emotion across 50 variables for 14 short films reported in the Annotation Study of the Emo-FilM dataset (Morgenroth et al. (2024)). Films were annotated continuously at a sampling rate of 1 Hz. 44 healthy volunteers produced annotations in real time during film viewing by moving a continuous bar on the side of the screen up and down a scale and were asked to make ratings according to their own emotional state using an adapted version of CARMA software (Girard (2014)). Participants annotated one item at a time. After extensive quality control a consensus annotation for each item and each film was calculated as the mean ratings from three to four raters for each item. We further processed this data based on the results of a factor analysis performed in (Morgenroth et al., 2023a). We computed continuous scores for each of three emotion dimensions ‘valence’, ‘arousal’ and ‘power’, sometimes termed ‘dominance’ (Russell and Mehrabian, 1977). Each emotion component was assigned to one of three dimensions based on the highest loading. We then summed the product of factor loading and emotion item for each dimension to achieve a time course for each emotion dimension. We furthermore applied smoothing and downsampling to match the temporal resolution of the fMRI data (TR=1.3 sec).

#### Individual difference scores

In the dataset, each subject in the fMRI study completed a series of questionnaires, which provides us 15 scores for each participant, resulting in 30 × 15 scores. These scores include the five subscales of the Big Five Inventory (Goldberg, 1990) (Extraversion, Agreeableness, Conscientiousness, Neuroticism, and Openness), the three subscales of the Depression Anxiety Stress Scales (Lovibond and Lovibond, 1995) (State Depression, State Anxiety and State Stress), four subscales of the Behavioural Activation and Behavioural Inhibition Scales (BIS/BAS)(Carver and White, 1994) (Drive, Fun-seeking, Reward responsiveness and Behavioural inhibition), two subscales of the Emotion Regulation Questionnaire (Gross and John, 2003) (Cognitive Reappraisal and Expressive Suppression), and a score of the emotional impact of the COVID pandemic, which was derived from a custom scale (Pandemic stress). Throughout, we term these 15 scores - individual difference scores.

### 2.2. FMRI Analysis Pipeline

An overview of our analysis pipeline is illustrated in Fig. 1. Briefly, we extract functional gradients from film and rest fMRI, and then evaluate inter-subject variability of these functional gradients as well as their potential to predict individual differences. We then explore an emotion-informed approach by selecting specific frames from film fMRI based on emotion rating scores to explore how experienced emotion modulates inter-subject variability and prediction of individual differences.

**Figure 1:**
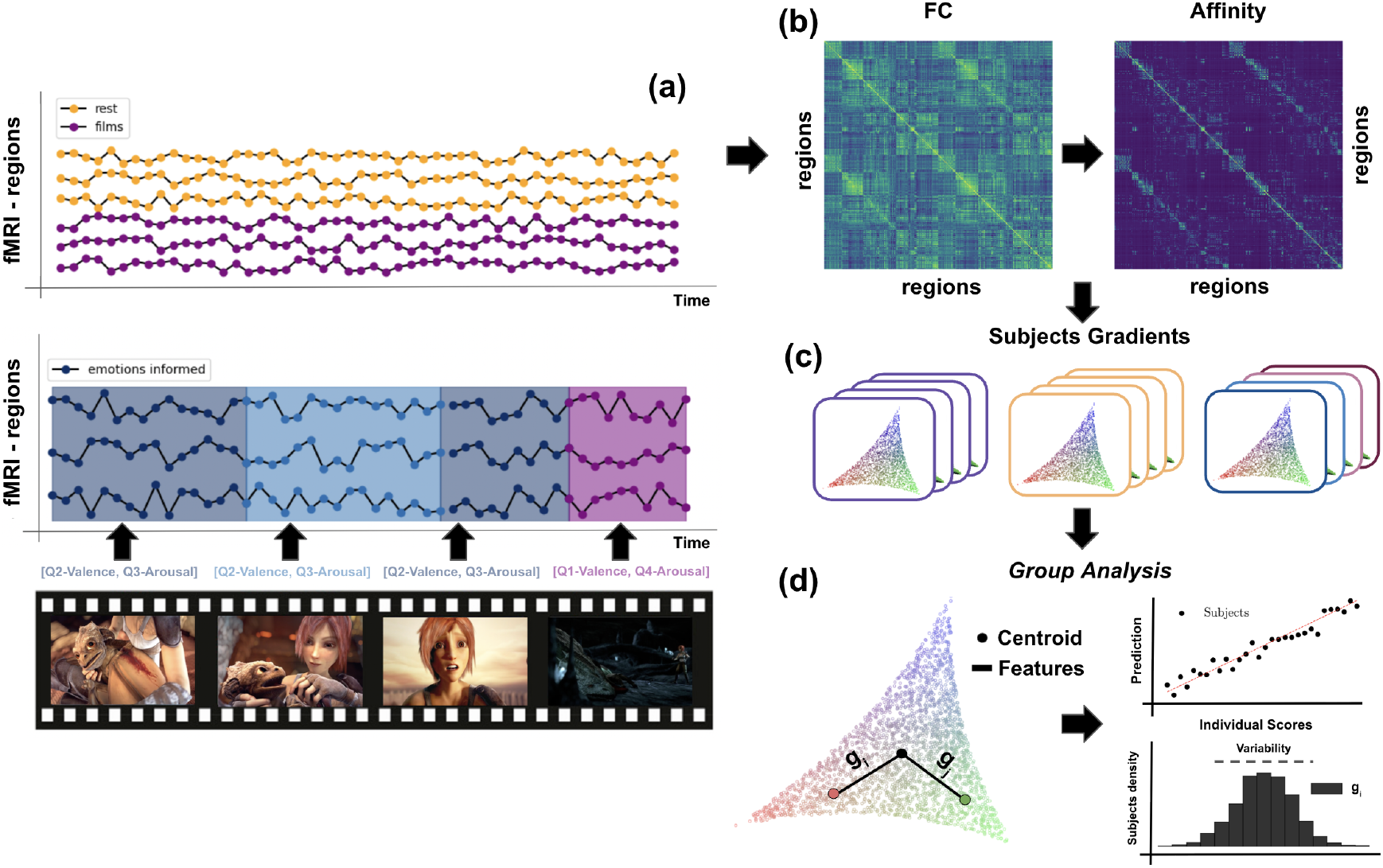
Study of rest, films and emotion-informed functional gradients in highlighting individual differences and variability. a) fMRI timecourses of films, rest, and segments of timecourses within films associated to quartiles of emotion scoring (Methods.2.2.4). b) FC matrices are calculated for each subject, as well as affinity matrices which are computed by Pearson correlation, from the row-thresholded FC matrix. c) Functional gradients are computed from each subject’s affinity matrix. (specific parameters are detailed in Methods.2.2.2). d) Gradients features (distance to centroid, Methods.2.2.2) are then used for both predictability (performing Lasso regression with respect to a given individual difference score) and computing inter-subject variability (standard deviation of gradients’ features) as detailed in Methods.(2.1, 2.2.6).

#### 2.2.1. Functional Connectivity

As a basis for computing functional gradients, we first built the FC matrix. To this end, we computed the pairwise correlations between all regional timecourses, resulting in an FC matrix of size 414×414. We computed both individual FC matrices and a group-level FC matrix constructed by concatenating all the z-scored individuals’ timecourses prior to applying pairwise correlation. Additionally, we computed a single FC matrix representative of the film condition by the concatenating the group timecourses across all films. The same procedure was applied for rest fMRI.

#### 2.2.2. Functional Gradients

These FC matrices were inputted into the BrainSpace Tool Box (Vos de Wael et al., 2020) to obtain functional gradients. Functional gradients represent a low-dimensional manifold of parcels onto which Euclidean distance between two points inversely relates to similarity between activity time courses of the corresponding two regions. To generate gradients at the individual and the group level, we first prune FC profile of each region (i.e., a specific row of the FC matrix) to 90%, sparsifying the very dense graph. Second, we compute an affinity matrix by correlating the newly generated rows pairwise, which symmetrizes the previously obtained pruned matrix. From the affinity matrix, we then project through diffusion embedding to a lower-dimension manifold following standard parameters α = 0.5, and an optimized number of diffusion time (Margulies et al., 2016). Choices regarding the construction of the gradients are mainly motivated by previous recommendations (Hong et al., 2020). Additionally, for gradients to be comparable, we employ BrainSpace’ implementation of Procrustes alignment that allows for scaling, rotation, and flipping of the gradients. More specifically, we use the generalized Procrustes method, which iteratively aligns all gradients to the mean across subjects. We align each type (rest, films, emotion-informed) of gradients within the set of subjects and when comparing between types we align using the group level rest gradients as reference. We extract features from FC gradients, such as the distance from a region’s embedding to the embeddings’ centroid and use these features to compare between FC gradients. This has several benefits over immediate gradient scores, gradients’ similarity metrics or gradients’ explained variance (Samara et al., 2022; Kong et al., 2023) since the distance to centroid is invariant to rotation and shifts it makes up for possible alignments errors. Moreover, it reduces feature space from three entries (one per gradient) to a single one, yet still allows for per region analysis. Finally, the distance to centroid can be loosely viewed as the average distance to all other regions, thus expressing how similar a region is to all others.

#### 2.2.3. Inter-Subject Variability

Given individual FC gradients features, we quantify the inter-subject variability to be the standard deviation (across subjects) computed for each dimension of the feature space (Figure.1d). Consequently, for a set of FC gradients, we obtain one standard deviation value for each region. We consider as well the global inter-subject variability of a set of FC gradients which is the average of region-level standard deviations. In the same fashion, we can average this inter-subject variability across subsets of regions; e.g, all regions within the same functional network, instead of all regions.

#### 2.2.4. Emotion-Informed Gradients

The novel element that we bring here is the idea of gradients informed by emotion experience. We define quartiles of the strength of emotion experience for each of three emotion dimensions ‘valence’, ‘arousal’ and ‘power’, and then select frames of the fMRI timecourse according to these quartiles. We concatenate the frames for each quartile and dimension and compute an FC matrix as the input to obtain emotion-informed gradients. Beyond the partitioning based on a single emotion dimension, we consider as well the intersection of quartiles across two emotion dimensions simultaneously. For example, we would select frames that fall within the lowest quartile (Q1) of valence, representing high negatively charged emotion, and to the second quartile (Q2) of arousal, representing moderate intensity. The resulting intersection [Q1-Valence, Q2-Arousal], would then be scenes associated moderately intense yet negatively charged emotion. Given that we use three different emotion dimensions, this leads to three pairwise combinations onto which we base our frame selection to derive the emotion-informed gradients. The percentiles are chosen across all films, which means that the sets of selected frames do not necessarily contain the same percentage of frames from each film run. While this process involves selecting frames, given a smoothed emotion experience timecourse, it effectively selects intervals of consecutive frames. This addresses the concern of capturing temporal dynamics of an emotion experience.

#### 2.2.5. Test-Retest Reliability for Films

The data from the 14 films are partitioned into two sets, as detailed in Supplementary Table 1, to provide a test-retest reliability assessment. We perform Intra-Class-Correlation (ICC) and discriminability analysis to determine how reproducible functional gradients are regardless of the film stimuli across subjects. For ICC, the classes would be subjects i.e verifying that a subject’s gradients are consistent between test, retest and the discriminability yields how often the retest’s gradient of a subject is the closest gradients to the test’s gradient of the same subject among all the other test gradients. Both metrics are computed for subsets of the test and retest sets to have an ICC and discriminability value for different input length. The procedure is repeated for the full films and the emotion-informed gradients. It validates the emotion-guided frame selection method we introduce here.

#### 2.2.6. Prediction of Individual Differences

With the goal of investigating how informative a given gradients is to a specific individual difference score, we employ a lasso regression on features of the gradients. The regression can be formulated as follows:

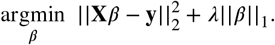

In summary, using the gradients’ features **X** (Methods.2.2.2), we predict the individual difference profile of each subject **y** (Methods.2.1). The model is fit on a train subset of 20 subjects and predictability is estimated with the remaining subjects. The prediction is evaluated on an outer fold test set while we use the outer fold training set to both optimize our Lasso regression weights *β* and L1 penalty parameter *λ*. The evaluation is reported as the Pearson correlation between predicted values on test set and original score. Since the outer fold choice influences the performance, we repeat the choice of outer fold a total of 250 times and generate a distribution of correlations. This scheme is used to compare between film gradients, emotion-informed gradients and rest gradients.

#### 2.2.7. Predictability and Variability Correlation

We applied Pearson correlation between predictability and inter-subject variability to understand a possible relationship.

##### Null model

The inter-subject variability values are shuffled across quartiles and once again we computed the correlation between inter-subject variability and predictability to generate one sample of the null distribution. We repeat this process 250 times to generate a null distribution of correlations. The tested correlation value is significant if it is either extremely negative (smallest 5%) or extremely positive (largest 5%).

### 2.3. Data and Code Availability

MRI and behavioural data as well as the film annotations are part of the EmoFilM dataset (Morgenroth et al., 2024). All code for the analyses and figure plotting used for this paper can be found following this link Github Repository.

## 3. Results

### 3.1. Gradients Layout in Films and Rest

Functional gradients for films and rest are displayed in Figure.2. Rest and films principal gradient are organized in a cortical unimodal-transmodal way with posterior, mainly visual areas anchored on one extreme and the frontal regions anchored on the other extreme. On the second gradient of both rest and films, we also find a unimodal-transmodal organization wherein we retrieve somatomotor regions on one end and regions from the default mode network (DMN) on the other end of the gradient. On the third gradient we see a stronger difference between rest and films. In films, the DMN and limbic (including medial and lateral temporal) regions constitute both ends of the third axis which brings about once again a unimodal-transmodal layout, while in rest, there is no clear anchoring. Together, the top 3 film-modulated cortical gradients are modality-specific, hierarchical gradients with unimodal to transmodal setup differing from rest gradients found in literature (Margulies et al., 2016). Note that on the first two gradients in both rest and films, subcortical regions are generally localized between the anchoring networks distant from the extremities, with a slight bias toward the transmodal type, while for the films’ third gradient specifically, different subcortical regions are localized on opposite extremes of the gradient with striatal and hippocampus areas strongly associated with the limbic pole.

**Figure 2:**
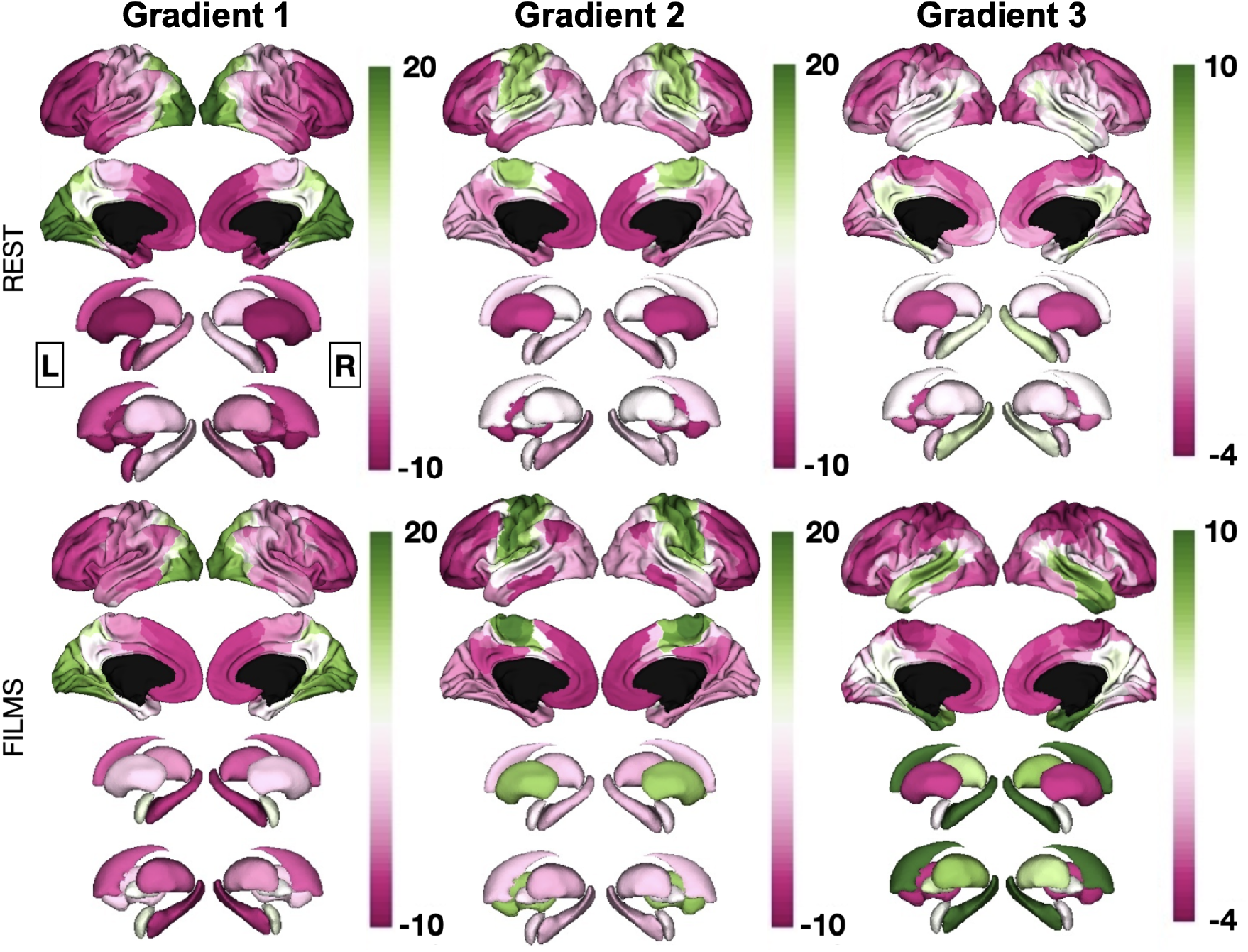
Group level functional gradients’ for rest and films on cortical and subcortical surfaces (legends in Supplementary Figure 9). The first 3 gradients are plotted for both films and rest with films gradients being procrustes aligned to rest gradients. Further comparative analysis on the films and rest gradients can be found in Supplementary Figure 2.

### 3.2. Inter-Subject Variability for Films and Rest

We computed the standard deviation of each region’s position within the gradient space across subjects for both rest and film. We find that this inter-subject variability is larger for rest than films gradients’ (Figure.3A). We further explored this pattern on the network level to better characterize this distribution. Only in visual, dorsal attention, somatomotor, and default-mode networks was the difference in inter-subject variability between rest and film significant (Figure.3B) (*p* = 0.0001). Notably, in contrast, we found that the somatomotor network is significantly more stable in rest than in films (*p* = 0.0001).

**Figure 3:**
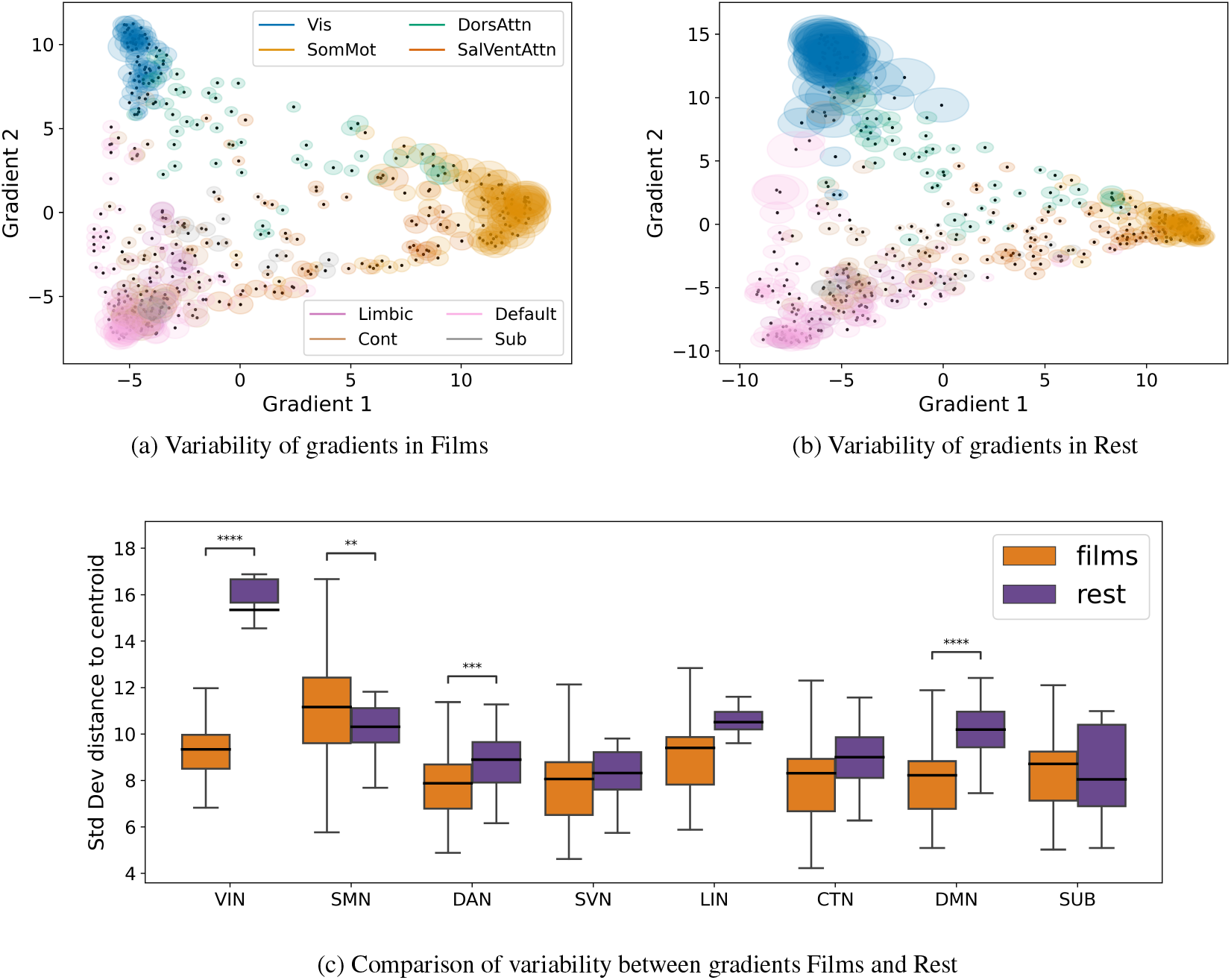
(a,b) Inter-subject variability (standard deviation) for different parcels for both gradients computed from films runs and gradients computed from rest runs. The first two gradients are shown here. (c) Distributions of standard deviation of each parcel’s distance to centroid (grouped per network). Unpaired t-test is used to compare between films and rest conditions due to sample difference (14 films versus 1 rest) and all numerical statistics are listed in Supplementary Table 2.

### 3.3. Predicting Individual Differences in Films and Rest

We find that using regression based on film gradients, we can predict individual differences in state anxiety. On the other hand, for rest (Figure.4), we find that we can predict scores of openness. State anxiety is statistically better predicted from films (*p* = 0.035) while openness is better predicted from rest (*p* = 0.001). No other scores’ pair-wise differences yielded statistical significance. Therefore, we focus onward on both state anxiety and openness score and look at their prediction using an emotion-informed approach.

**Figure 4:**
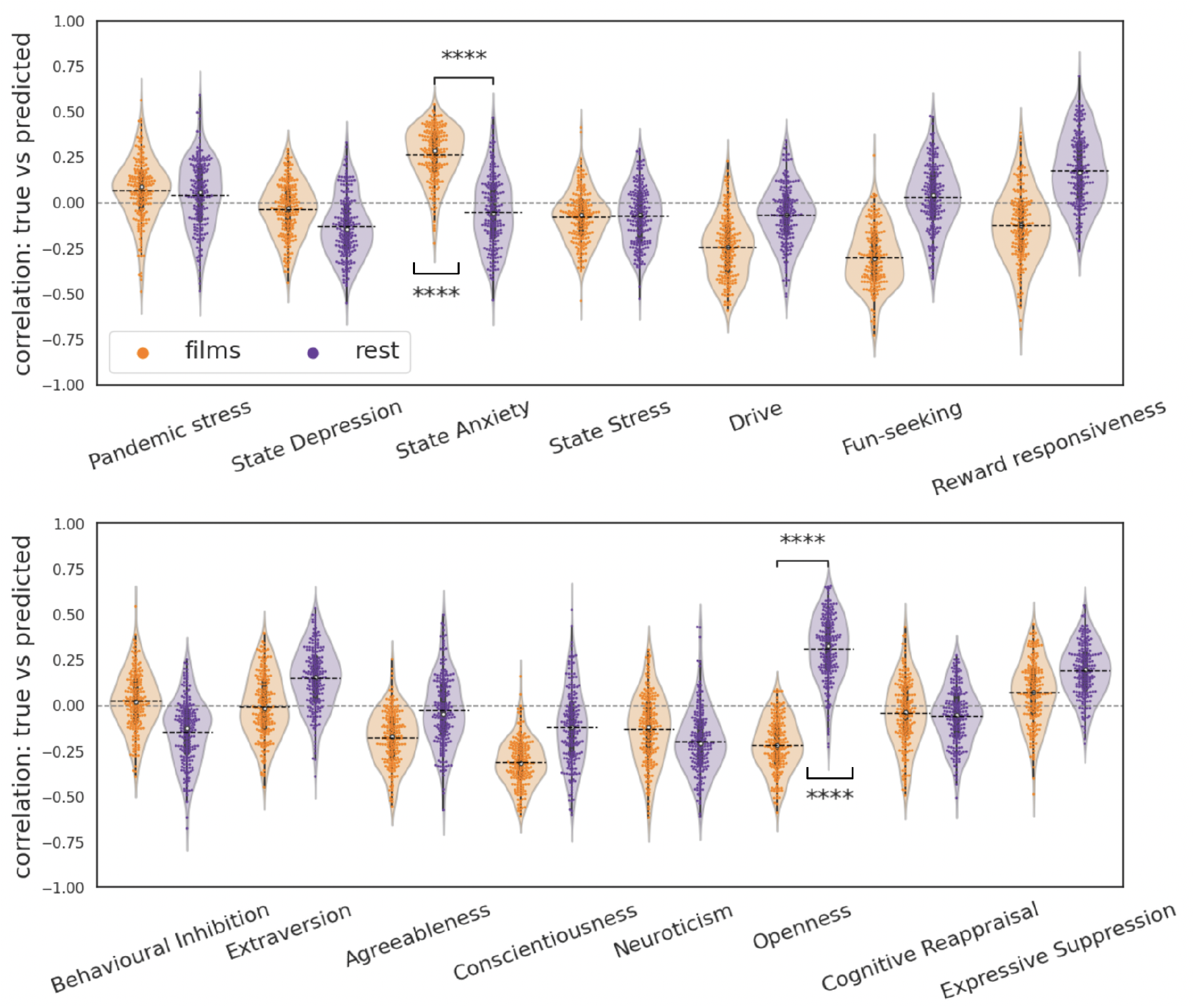
Prediction results (correlation with consensus annotation) on the various test folds for all individual differences in the dataset. The individual differences are introduced and detailed in Methods.2.1. Two-sample t-tests in concordance with Cliff’s *δ* with large effect threshold of 0.474 (considering sample size) was employed. Pairs of conditions are similarly tested. Statistically significant fold distributions are shown in an underlying bracket below the swarmplot while differences are shown in a second overarching bracket above the corresponding swarmplots. The numerical statistics are written in Supplementary Table 4.

### 3.4. Gradients Layout in Emotion-Informed Gradients

Gradients based on selected timepoints according to emotion annotations were found to organize in a layout similar to films gradients when considering group level gradients, as illustrated in Figure.5 with the gradients from Q4-Arousal.

**Figure 5:**
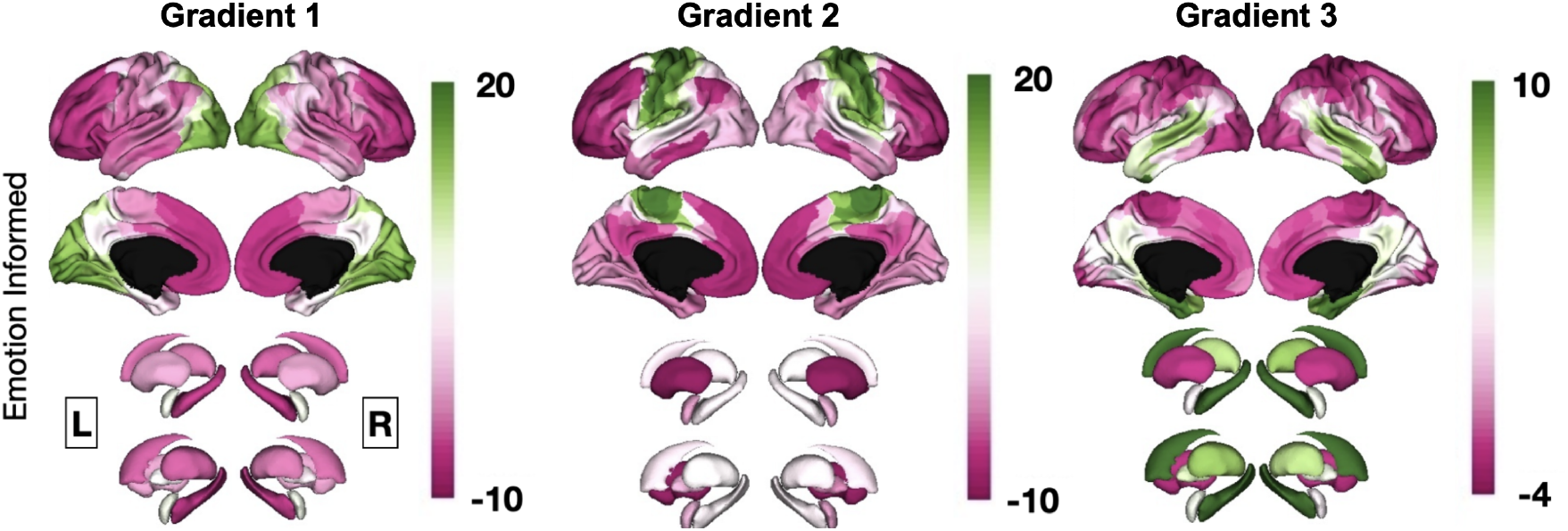
Group level functional gradients’ for emotion-informed gradients on cortical and subcortical surfaces. The first 3 gradients are plotted for emotion-informed gradients procrustes aligned to rest gradients in Figure.2. The emotion-informed gradients we show is that of Q4-Arousal. The same emotion-informed gradients is used in the following plots. Further comparative analysis on the emotion-informed gradients against films and rest gradients can be found in Supplementary Figure 3.

In Supplementary Figures, we show the remaining gradients from frame-selected time courses, which show very similar organizations. In this respect subsets of frames from films (if enough timepoints are selected) induce similar functional organization as the entire set of films. Consequently we observe a hierarchical layout anchored by the same sets of networks for emotion-informed gradients as in full films gradients.

### 3.5. Test-Retest Reliability for Emotion-Informed Gradients

In our test-retest, the test and retest sets are composed of different films altogether, the results, yet, show that individual films gradients remain stable (mean ICC in films = 0.88 at full split; mean ICC in Frame Selected = 0.74 at the size of a quartile). We also observe that full films gradients have a higher ICC score compared to emotion-informed gradients thanks to the higher number of frames averaging variability of individual frames. Cortical distributions of ICC scores at half split show frontal lobes to have lowest ICC scores across all quartiles (Figure.6C). As for Discriminability, we use all 30 subjects at varying length for the test-retest sets. From the Discriminability scores, we observe strong test-retest reliability for all quartile ranges of emotion-informed gradients and as well for full films. At half split (7 films included), emotion-informed gradients’ Discriminability reaches DIS = 0.97 while full films gradients’ Discriminability reaches DIS = 1.0. In emotion-informed gradients, the choice of partitioning the timecourses into quartiles is made considering the number of frames obtained (Supplementary Figure 4) when intersecting quartiles. ICC and Discriminability scores for other partitioning size are shown in Supplementary Figure 5 and 6. Ultimately, we find that emotion-informed gradients for all quartile ranges possess strong test-retest reliability, showing that emotion-informed gradients computed from frames associated with the same interval of emotion experience are similar regardless of the film which evokes the experience.

**Figure 6:**
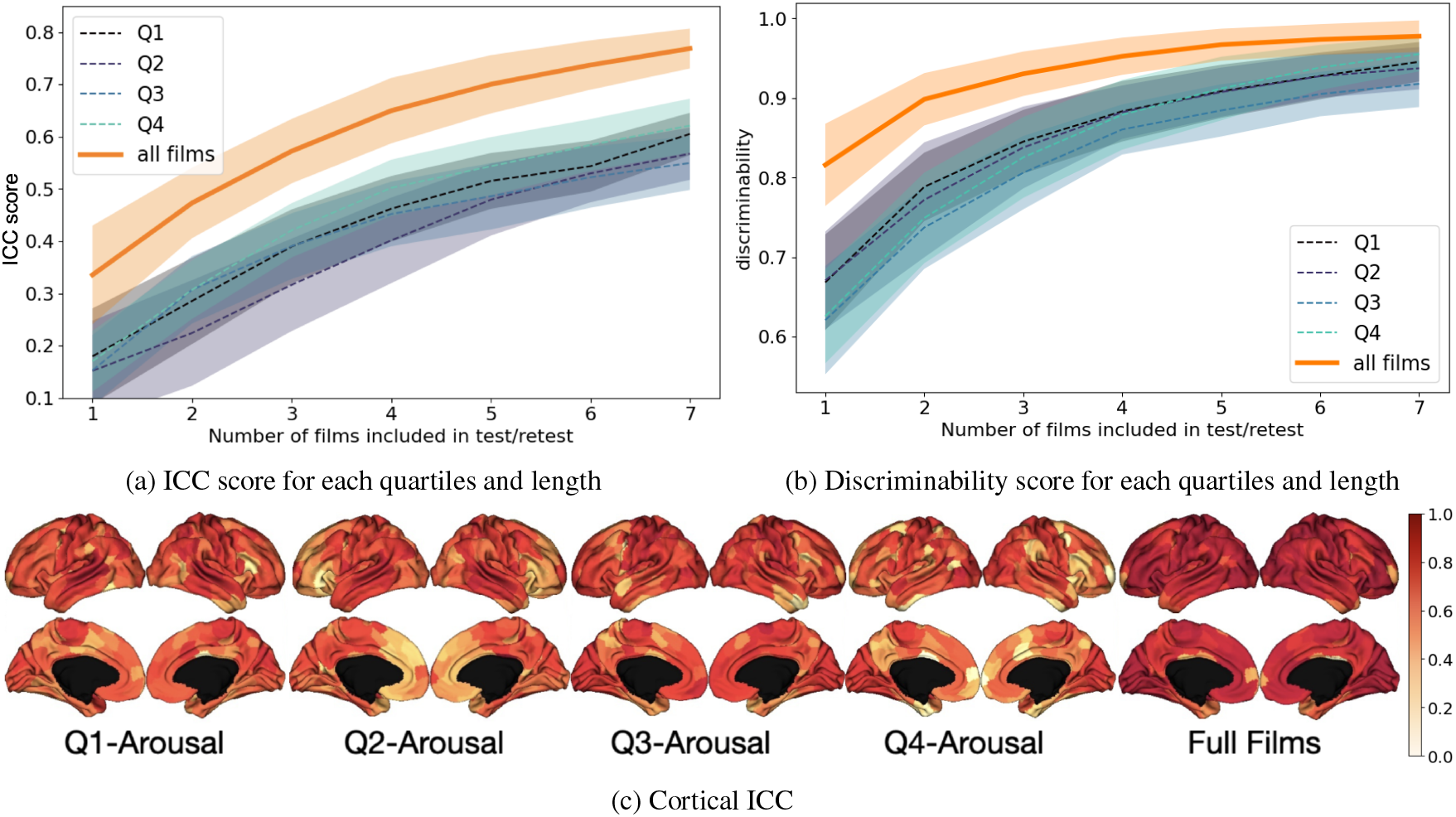
(a) Averaged ICC score over all parcels (single rater, random trial and absolute agreement). The ICC scores are computed for different size of test-retests and over full films gradients’, Q1 to Q4 emotion-informed gradients. (c) The ICC scores of each parcel are shown on cortical surface for few emotion-informed gradients. (b) Averaged Discriminability score over all parcels. Discriminability is computed for different size of test-retests and over full films gradients’, Q1 to Q4 emotion-informed gradients.

### 3.6. Inter-Subject Variability in Emotion-Informed Gradients

We inspect the inter-subject variability of functional gradients derived from the intersection between two different emotion scores’ quartile and thus we have 16 distributions of standard deviations for each of the 3 pairs of scores. Among some notable trend is the decreasing variability as arousal increases whereas variability increases as power increases (Figure.7A). In Supplementary Figure 4, we report the number of frames for each intersection between pairs of different emotion scores’ quartiles and find that generally an acceptable number of frames is yielded.

**Figure 7:**
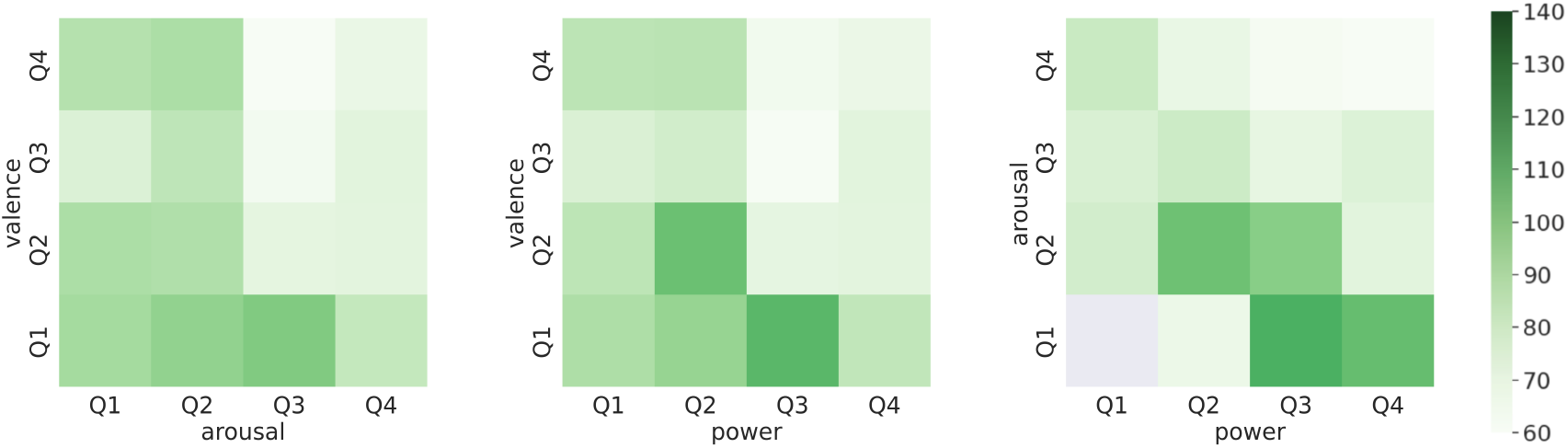
Inter-subject-variability of emotion-informed gradients. The variability is unit-less as it is computed from gradients’ scores (Methods.2.2.3) and emotion-informed gradients are computed from intersection of percentiles Supplementary Figure 4.

### 3.7. Predicting Individual Differences in Emotion-Informed Gradients

Results of nested folds cross validation with Lasso yielded scores for pairs of quartiles (Valence/Arousal, Valence/Power, Arousal/Power). The two highest correlation intersections of quartiles are [Q2-Valence, Q3-Arousal] and [Q3-Power, Q3-Arousal] for state anxiety as shown in Figure.8A. Then the two highest correlation intersections of quartiles are [Q1-Valence, Q1-Arousal] and [Q3-Power, Q1-Arousal] for openness as shown in Figure.8B (with statistical significance in prediction shown in the p-value table in Supplementary Figure 7). [Q2-Valence, Q3-Arousal] are frames of negative valence and medium high arousal, similarly [Q3-power, Q3-Arousal] are frames of high power and high arousal for state anxiety. Now for openness, [Q1-Valence, Q1-Arousal] are of negative valence and low arousal and [Q3-Power, Q1-Arousal] are frames of high power and low arousal. In general, state anxiety is better predicted in high arousal inducing frames while openness is better predicted in low arousal inducing frames. Comparing prediction we obtain a maximal prediction score of *ρ*_emo-info_ = 0.33 averaged on test folds while originally on full films we have *ρ*_full_ = 0.26 and on rest we have *ρ*_rest_ = 0.03 for state anxiety. For openness, we obtain a maximal prediction score of *ρ*_emo-info_ = 0.32 averaged on test folds while originally on full films we have *ρ*_full_ = 0.03 and on rest we have *ρ*_rest_ = 0.21.

**Figure 8:**
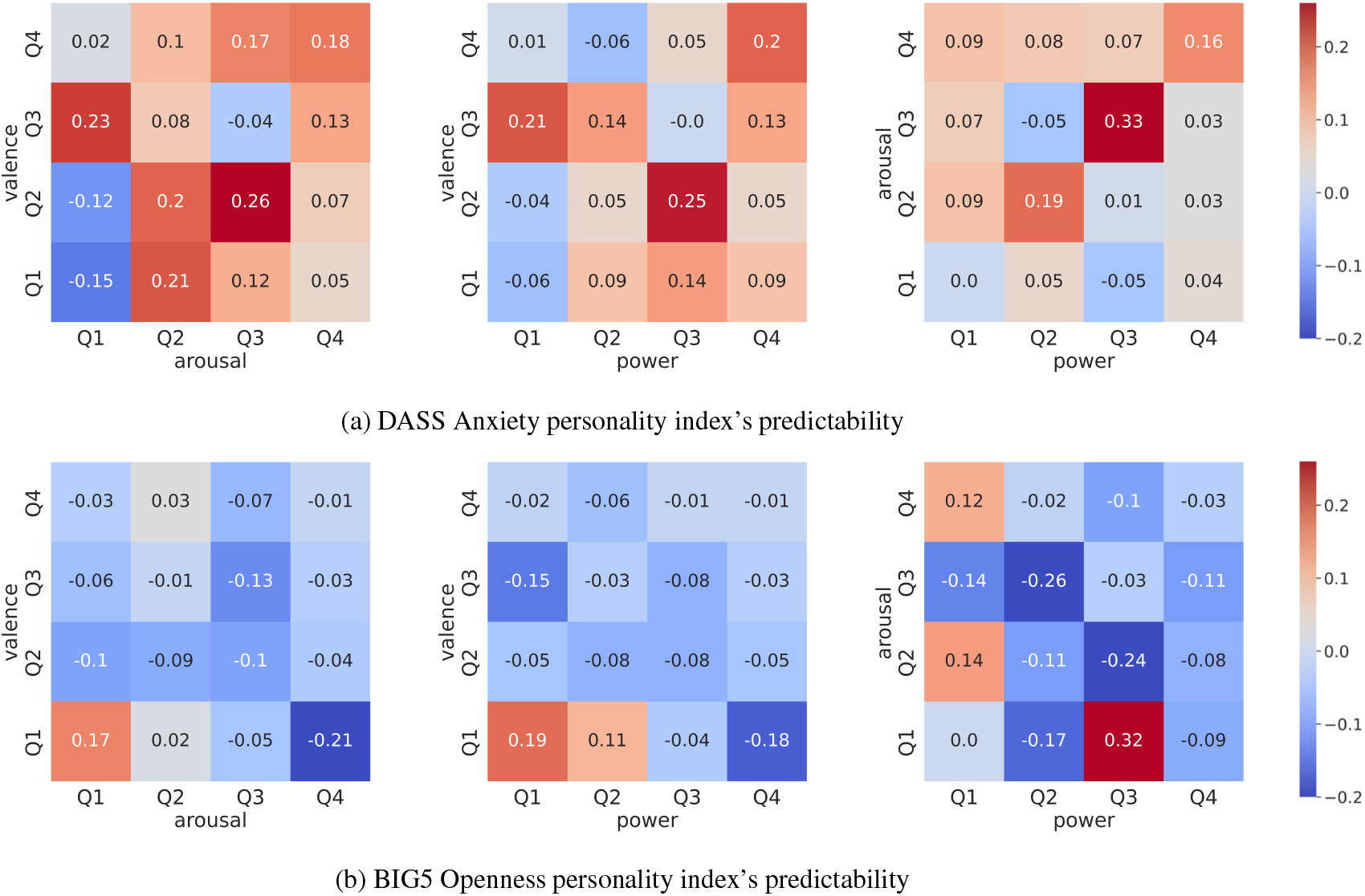
(a) Prediction results (correlation with consensus annotation) on the various test folds for individual difference: **state anxiety**. Results are computed using as feature vectors all different combination of frame selections and standard deviations of results on the set of folds are detailed in Supplementary Figure 8. (b) Prediction results (correlation with consensus annotation) on the various test folds for individual difference: **openness**. Results are computed using as feature vectors all different combination of frame selections and standard deviations of results on the set of folds are detailed in Supplementary Figure 8.

### 3.8. Predictability and Variability in Films

In the previous section we showed that certain frames within films performed better in the identification of individual differences. Therefore, we related inter-subject variability of those frames with their power to predict individual differences. Specifically, regarding the predictability of state anxiety as seen in Table 2, we find a significant negative correlation between predictability and variability of Limbic network and Visual network. On the contrary, for openness we find a significant positive relationship between inter-subject variability of the Visual network and predictability (Table 2). We further tested for significant differences of this relationship between both scores. This confirmed, that the prediction of state anxiety is modulated by inter-subject variability to a greater degree than openness. Specifically, state anxiety is better predicted by frames with low inter-subject variability while openness is better predicted by frames with higher inter-subject variability.

**Table 1.** Two-way ANOVA with two independent factors being condition (rest-films) and networks. The significance of the difference of means is shown and interaction between condition and networks as well.

|  | df | sum_sq | mean_sq | F | PR(>F) |
| --- | --- | --- | --- | --- | --- |
| C(cond) | 1 | $0.318 \times 10^6$ | $0.318 \times 10^6$ | 122.74 | 0.0 |
| C(network) | 7 | $2.95 \times 10^6$ | $0.422 \times 10^6$ | 163.11 | 0.0 |
| C(cond):C(network) | 7 | $1.20 \times 10^6$ | $0.171 \times 10^6$ | 66.00 | 0.0 |
| Residual | 6194 | $16.026 \times 10^6$ | 2587.40 | NaN | NaN |

**Table 2.**
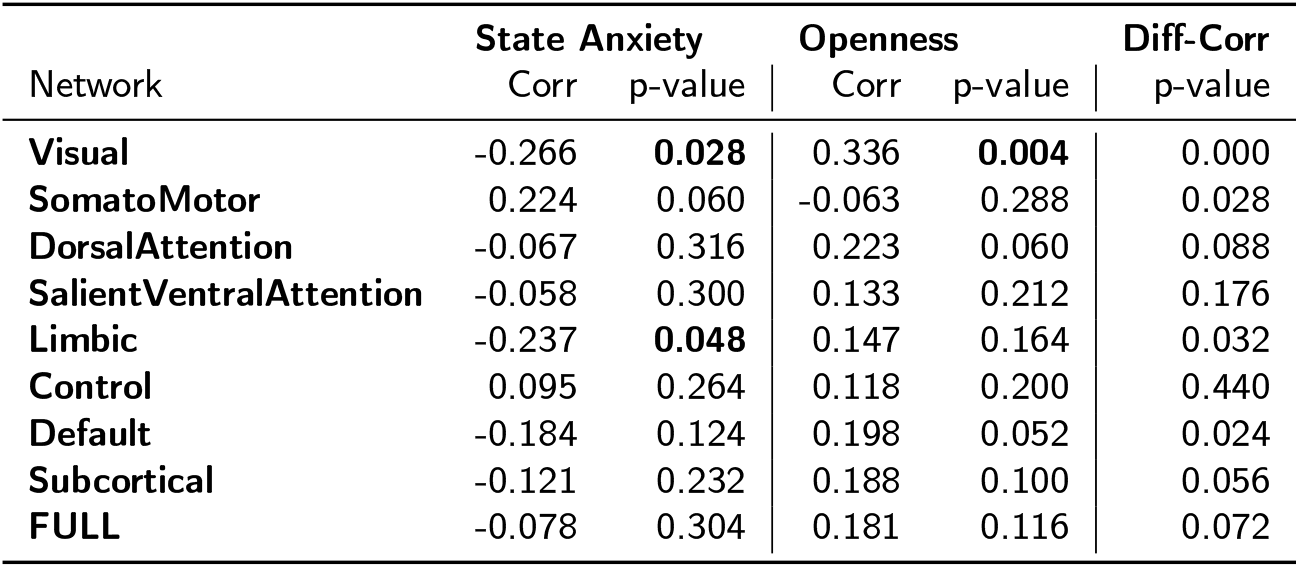
Correlation between mean variability per percentile against predictability scores on **state anxiety, openness** and statistics on the two score’s differences. The statistics are computed from non-parametric testing and the partitioning is based on intersection over quartiles from different pairs of emotion dimension.

| Network | State Anxiety |  | Openness |  | Diff-Corr |
| --- | --- | --- | --- | --- | --- |
|  | Corr | p-value | Corr | p-value | p-value |
| **Visual** | -0.266 | **0.028** | 0.336 | **0.004** | 0.000 |
| **SomatoMotor** | 0.224 | 0.060 | -0.063 | 0.288 | 0.028 |
| **DorsalAttention** | -0.067 | 0.316 | 0.223 | 0.060 | 0.088 |
| **SalientVentralAttention** | -0.058 | 0.300 | 0.133 | 0.212 | 0.176 |
| **Limbic** | -0.237 | **0.048** | 0.147 | 0.164 | 0.032 |
| **Control** | 0.095 | 0.264 | 0.118 | 0.200 | 0.440 |
| **Default** | -0.184 | 0.124 | 0.198 | 0.052 | 0.024 |
| **Subcortical** | -0.121 | 0.232 | 0.188 | 0.100 | 0.056 |
| **FULL** | -0.078 | 0.304 | 0.181 | 0.116 | 0.072 |

## 4. Discussion

The aim of this study is to investigate how affective processes impact FC reorganization, considering inter-subject variability and subject individual difference prediction using functional gradients. In this study, we sought to understand how affective processes impact the cortico-subcortical organization of functional connectivity with regards to inter-subject variability and prediction of individual differences during rest and film fMRI. To this end we employed FC ICC score for each quartiles and length (b) Discriminability score for each quartiles and length gradients and further expanded their use in an emotion-informed approach. We selected frames in film fMRI based on emotion experience to better understand the effect of current emotion state on gradient organization.

We reproduced a similar macrostructural cortical organization along the first three gradients as reported in previous work (Margulies et al., 2016; Samara et al., 2022). Additionally, we show that variance is distributed more evenly across the first three gradients in film fMRI than in rest and that a differential organization within subcortical regions is especially apparent along the third gradient. Accordingly, unlike some prior work, we decided to include the third gradient in our analysis.

When investigating the inter-subject variability of these gradients, we found it to be considerably lower during films than during rest as expected. We believe this is due to high levels of inter-subject synchronization by film content (Hasson et al., 2004). Brain activation synchronizes between subjects when they experience the same time locked stimulation.

We were particularly interested in comparing prediction of individual differences from gradients in both rest and film. Sufficient variance between subjects is an important condition to be able to extract subject specific information, thus we wanted to understand how rest and film differed in prediction of individual differences given this reduction of variability during film fMRI.

Previous results with film fMRI (Finn, 2021; Finn and Bandettini, 2021) reported superiority of films in predicting individual differences in this affective domain. While, we found similarly that functional gradients from film fMRI are better predictors, it was specifically for state anxiety such that despite lower variance, the film-based prediction outperformed the rest-based prediction. And to our surprise our analysis demonstrates that films and rest may be differentially sensitive for the prediction of specific characteristics (such that here openness was better predicted by rest gradients). This is contrary to existing literature that shows task and film surpass rest fMRI for individual difference prediction (Finn and Bandettini, 2021). Therefore, more subtleties are left to investigate, one of which being, to further understand the effect of the very diverse emotion profiles of each film detailed in (Morgenroth et al., 2024) EmoFilm dataset.

Since inter-subject synchronization of brain activity during film watching (Li et al., 2021; Gruskin et al., 2020; Trost et al., 2015) has been shown to be dependent on the emotional content of the stimulus material, an emotion guided approach may help to better understand which features improve prediction of individual differences from film fMRI. We therefore also extracted emotion-informed functional gradients based on emotion experience. Emotion-informed gradients showed a similar macrostructure as gradients based on the whole time series, retaining the hierarchical relation between transmodal and unimodal networks. We further identified the minimum of frames needed for a desired reliability of emotion-informed functional gradients through a test-retest reliability. Results on inter-subject variability for emotion-informed gradients show decreasing variability with increasing arousal, indicating that synchronization is enhanced during events with stronger emotional intensity. This consolidates previous findings as to which features of films constrain inter-subject variability and justifies the interest of emotion-informed gradients based on emotion experience.

We found significant differences in the prediction of state anxiety and openness between rest and films gradients. Therefore, we examined the effect of frame selection on the prediction of these metrics. For both scores we find that specific frame selections from the films possess stronger predictive power than others. Specifically for state anxiety, frames associated with negative valence and medium high arousal (i.e., stronger negative content) yielded the highest predictions meaning that frames that are likely to convey threat allow for best prediction of individual differences with regards to anxiety. This suggests that emotion-informed gradients allow to target distinct aspects of individual differences and extract more informative frames. From this perspective, it may even be possible to associate specific features of emotion experience to a given individual difference, further enhancing idiosyncrasy. In contrast, prediction of openness overall was best with rest fMRI and similar predictability was only obtained from frames of low arousal. Finally, when relating predictability to inter-subject variability, we show that these two variables are negatively correlated for state anxiety in Limbic and Visual network, while they are positively correlated in Visual network for openness. Overall, functional gradients that have low intersubject variability and that potentially reflect very specific emotional content bring about stronger predictive power for state anxiety, whereas a less constrained stimulus condition produces more informative functional gradients concerning openness.

In this work, we had a sample size of 30 subjects and thus, despite the large amount of data per subject, this first limited the prediction of individual differences when using regression models and also the quantification of gradients’ variability across subjects. Test-retest reliability was also computed across our set of 30 subjects. As a next step it is important to extend this analysis to a larger dataset. In the future, it will be of interest to extend this method to a greater variety of measures beyond personality and emotional state, for example performance measures from different cognitive tasks or phenotypic features, and thus uncover both their shared and distinctive underlying dimensions. Additionally, our method of frame selection may be extended beyond emotion dimensions and be computed according to other metrics, such as physiological measurements during rest or specific experimental blocks in task fMRI. Our analysis further highlights advantages of films over rest as found previously (Finn and Bandettini, 2021; Vanderwal et al., 2017) yet also disadvantages. Conversely, the choice of film or rest fMRI may be a question of which features would benefit from a more constrained stimulation and which come to light more reliably during unconstrained activation. This may also inform work related to brain fingerprinting (Van De Ville et al., 2021; Griffa et al., 2022), for example posing the question of whether there is a modality specific brain fingerprint. Finally, a possible generalization of our approach employing fully dynamic functional gradients based on dynamic functional connectivity may be a promising avenue to obtain a continuous range of predictive power along a continuous range of inter-subject variability.

## Conclusion

We investigated functional gradients for film and rest fMRI and introduced emotion-informed functional gradients for the film data. We first show differences in rest and film regarding inter-subject variability and prediction of individual differences. In an effort to better understand which elements of film contribute to advantages in predicting individual differences we then apply frame selection based on emotion experience to extract emotion-informed gradients. We find that film and rest highlight unique features among individual traits and may therefore be exploited for differential uses depending on one’s variables of interest. Intriguingly despite its constrained inter-subject variability due to stronger inter-subject synchronization, gradients from film fMRI outperform rest for some features of individual differences and not others. In the context of the large interest and the use of naturalistic stimuli in fMRI, this study underscores the importance of considering film content with regards to emotion experience, but also confirms the important position of rest fMRI as a paradigm in neuroscience.

## Acknowledgment

All authors declare that they have no conflict of interest. This work was supported by the Swiss National Science Foundation Sinergia grant CRSII5_180319. We thank the staff of the imaging platform at the Brain and Behavior Lab (BBL) for technical support and advice during data collection as well as Stefano Moia, Raphael Fournier, Michal Muszynski, Maria Ploumitsakou and Marina Almató-Bellavista for their contribution to the dataset used in this paper.

## Author Contributions

MC: Methodology (equal), Visualization (lead), Formal Analysis (lead), Writing – Original Draft Preparation (equal), Writing – Review & Editing (equal) LV: Formal Analysis (supporting), Methodology (equal), Writing – Review & Editing (equal) PV: Funding Acquisition (equal), Supervision (supporting), Writing – Review & Editing (equal) DvdV: Funding Acquisition(equal), Conceptualization (equal), Supervision (lead), Writing – Review & Editing (equal) EM: Conceptualization (equal), Methodology (equal), Visualization (supporting), Writing – Original Draft Preparation (equal), Writing – Review & Editing (equal), Supervision (lead)

## Compliance with Ethical Standards

Conflict of Interest: The authors declare that they have no conflicts of interest.

## Notes

### Competing Interest Statement

The authors have declared no competing interest.

