## Supplementary for "Individual differences of cortical and subcortical emotion-informed functional gradients"

### Supplementary Figures

- List of films used in the dataset (Table.1)
- Difference films-rest along gradients (Figure.1)
- Comparisons between films and rest gradients (Figure.2)
- Comparisons between emotion informed gradients and films and rest gradients (Figure.3)
- Number of frames composing each emotion-informed gradients (Figure.4)
- Statistics on intersubject variability films-rest (tables statistics of pair values Table.2)
- Statistics of prediction of individual scores (Table.3)
- Statistics of paired differences all table (Table.4)
- Other percentile sizes for test retest (Figure.5, Figure.6)
- The standard deviation of test folds (Figure.7)
- Test statistics of original distribution and null distribution for each intersections (Figure.8)

**Table 1**
List of films in the fMRI Emo-film dataset Morgenroth et al. (2024).

| **Film** | Duration (minutes:seconds) | Scan Duration (TR) | Genre | Test/Retest |
|---|---|---|---|---|
| After The Rain | 8:16 | 534 | Drama | Test |
| Between Viewings | 13:28 | 776 | Comedy/Drama | Test |
| Big Buck Bunny | 8:10 | 528 | Animation / Comedy | Retest |
| Chatter | 6:45 | 464 | Thriller | Test |
| First Bite | 9:59 | 613 | Romance | Retest |
| Lesson Learned | 11:07 | 665 | Drama | Test |
| Payload | 16:48 | 928 | Drama / Sci-Fi | Test |
| Sintel | 12:02 | 710 | Animation/ Fantasy | Retest |
| Spaceman | 13:25 | 744 | Romance/Drama | Retest |
| Superhero | 17:08 | 945 | Drama | Test |
| Tears Of Steel | 9:48 | 607 | Action/Sci-Fi | Retest |
| The Secret Number | 13.04 | 757 | Drama | Test |
| To Claire From Sonny | 6:42 | 460 | Drama | Test |
| You Again | 13:18 | 768 | Romance/Drama | Retest |

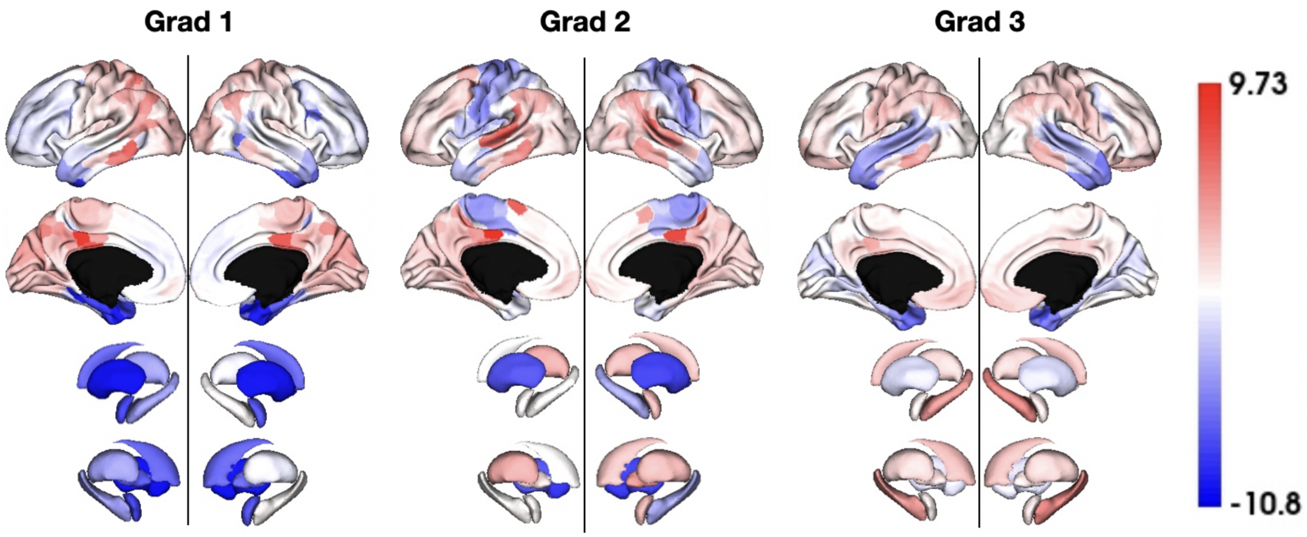


**Figure 1:** Distance between films and rest functional gradients along all gradients

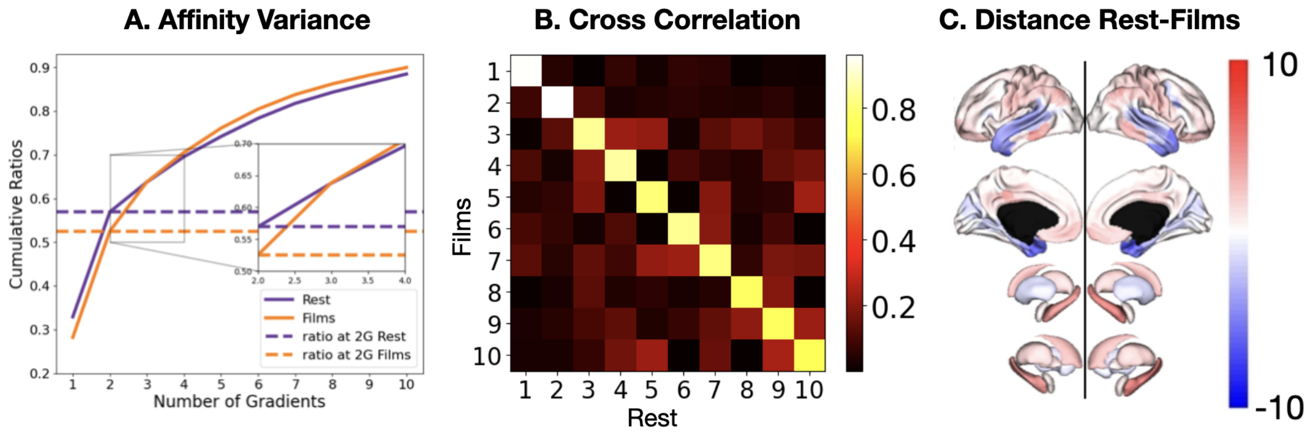


**Figure 2:** (**A**) Cumulative explained variance for the affinity matrix (proxy to original FC matrix) as a function the number of gradients added for both rest and films. (**B**) Correlation of films gradients and rest gradients scores. (**C**) Difference between films and rest gradients on the third gradients. (results for all gradients are shown on 1)

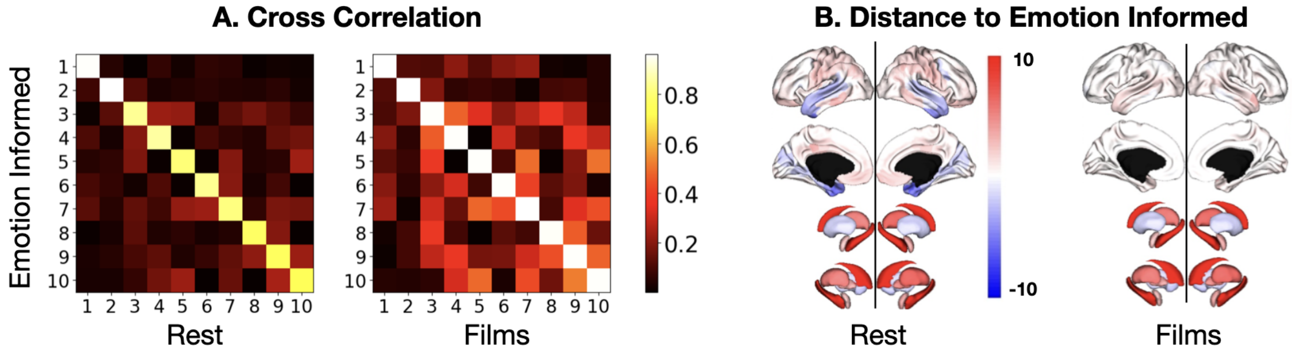


**Figure 3:** (**A**) Correlation of emotion-informed gradients and rest/films gradients. (**B**) Difference between emotion-informed gradients and rest/films gradients on the third gradients.

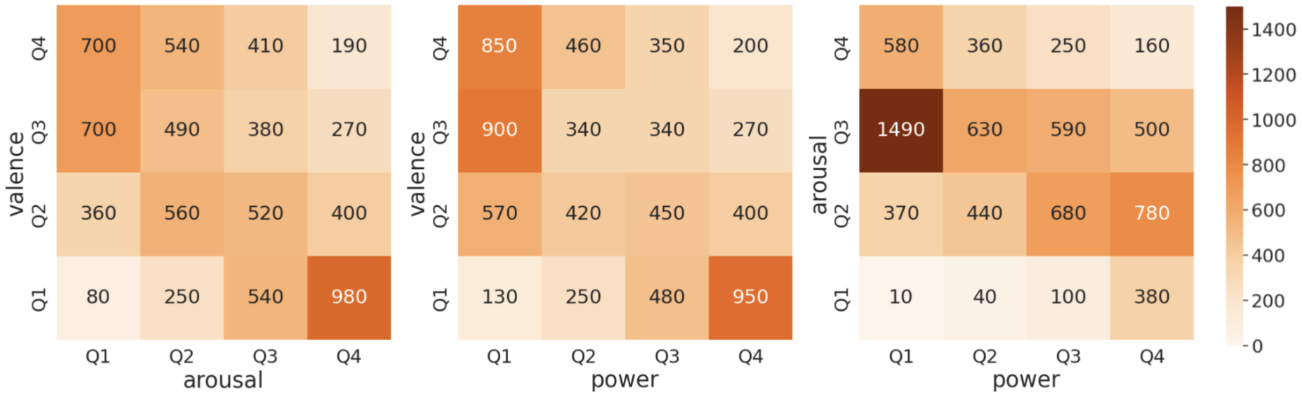


**Figure 4:** Number of frames used to compute different emotion-informed gradients.

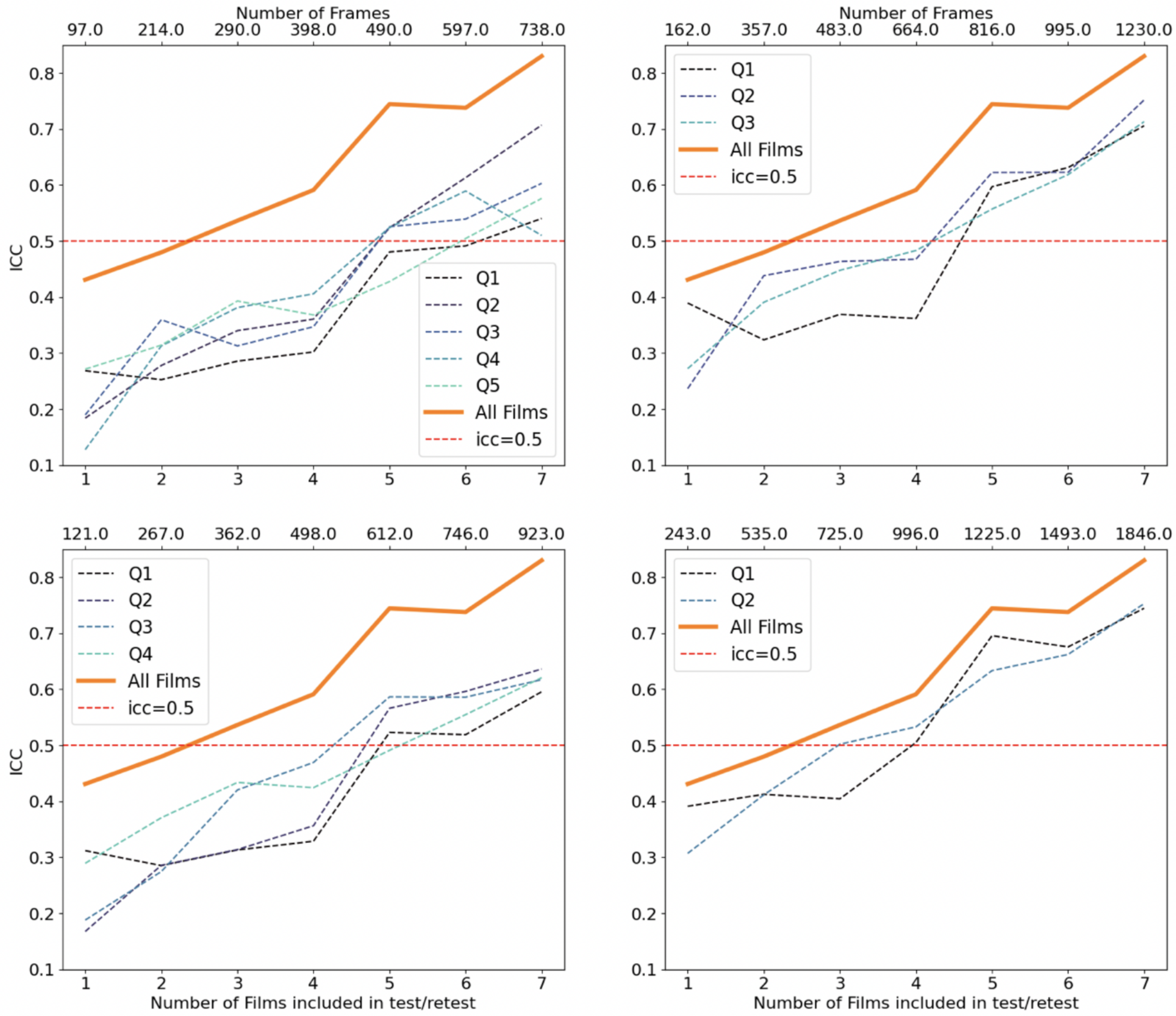


**Figure 5:** All tested percentiles in test-retest on ICC

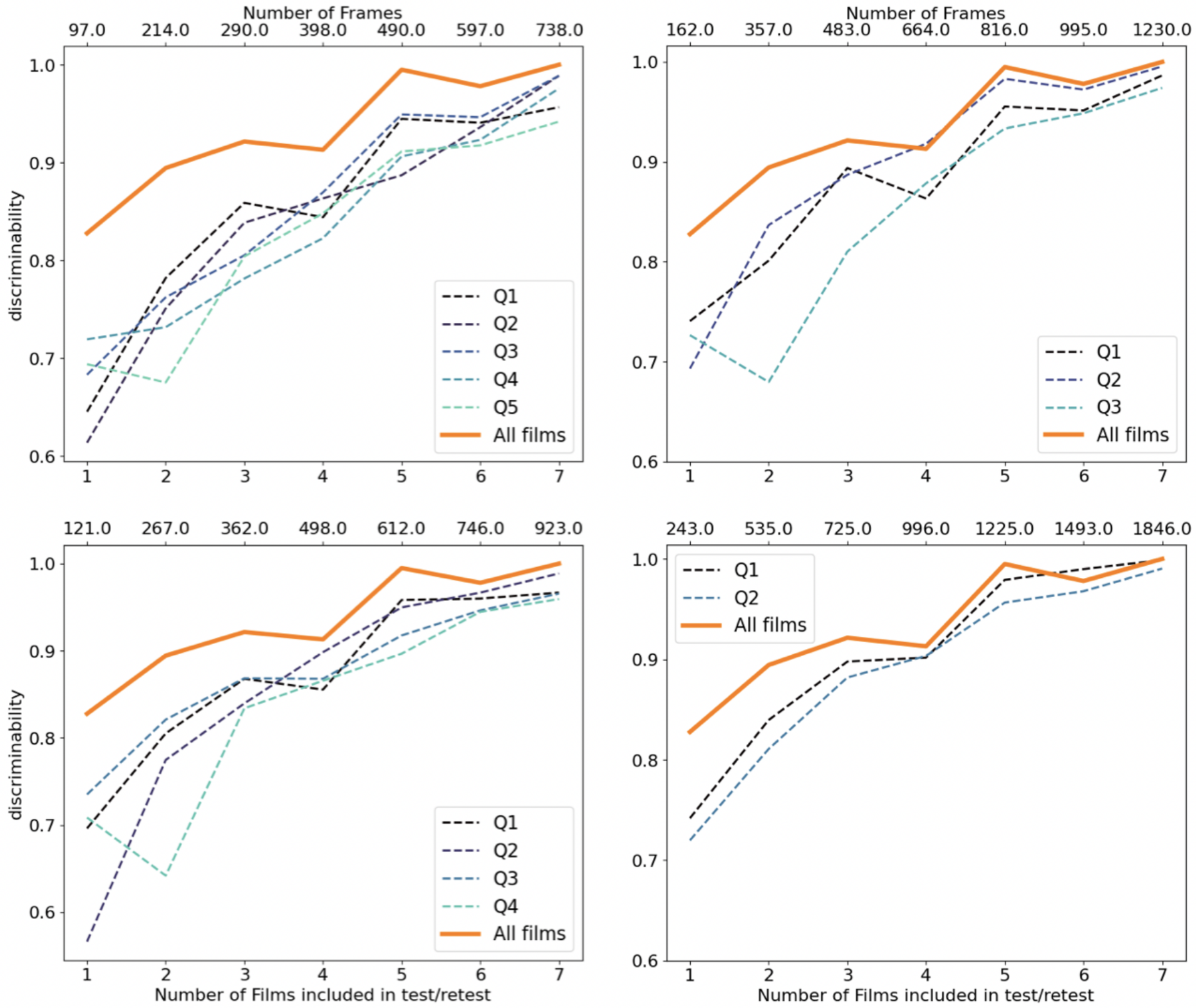


**Figure 6:** All tested percentiles in test-retest on Discriminability

**Table 2**
Wilcoxon test between variability in Films vs Rest on each networks

| Networks | Vis | SomMot | DorsAttn | SalVentAttn | Limbic | Cont | Default | Sub |
|---|---|---|---|---|---|---|---|---|
| pval | 0 | 0.0005 | 0.0011 | 0.9424 | 0.2919 | 0.4133 | 0 | 0.5125 |

**Table 3**
Two-sample t-test pvalues and Cliff's $\delta$ for rest and films gradients fold correlations.

| films | cov | dep | anx | str | bas_d | bas_f | bas_r |
|---|---|---|---|---|---|---|---|
| (pval, $\delta$) | (0.0001, 0.2155) | (0.9741, 0.1124) | (0.0, 0.7806) | (1.0, 0.324) | (1.0, 0.6513) | (1.0, 0.8051) | (1.0, 0.3766) |

| rest | cov | dep | anx | str | bas_d | bas_f | bas_r |
|---|---|---|---|---|---|---|---|
| (pval, $\delta$) | (0.0734, 0.0839) | (1.0, 0.492) | (0.9981, 0.1676) | (1.0, 0.2658) | (0.9997, 0.2039) | (0.011, 0.1358) | (0.0, 0.4154) |

| rest | bis | ext | agr | con | neu | ope | cr | es |
|---|---|---|---|---|---|---|---|---|
| (pval, $\delta$) | (1.0, 0.4704) | (0.0, 0.475) | (0.9849, 0.1252) | (1.0, 0.3192) | (1.0, 0.5809) | (0.0, 0.7517) | (0.9907, 0.1374) | (0.0, 0.6218) |

| films | bis | ext | agr | con | neu | ope | cr | es |
|---|---|---|---|---|---|---|---|---|
| (pval, $\delta$) | (0.4444, 0.0082) | (0.8431, 0.0582) | (1.0, 0.6454) | (1.0, 0.9336) | (1.0, 0.3145) | (1.0, 0.6902) | (0.9912, 0.1386) | (0.0012, 0.1759) |

**Table 4**
Two-sample t-test pvalues and Cliff's $\delta$ for paired differences between Rest and Films test fold correlations.

| paired | cov | dep | anx | str | bas_d | bas_f | bas_r |
|---|---|---|---|---|---|---|---|
| (pval, $\delta$) | (0.2043, 0.0734) | (0.0, 0.3419) | (0.0, 0.7904) | (0.7303, 0.02) | (0.0, 0.5401) | (0.0, 0.8191) | (0.0, 0.7248) |

| paired | bis | ext | agr | con | neu | ope | cr | es |
|---|---|---|---|---|---|---|---|---|
| (pval, $\delta$) | (0.0, 0.5127) | (0.0, 0.462) | (0.0, 0.4468) | (0.0, 0.5495) | (0.0001, 0.228) | (0.0, 0.971) | (0.4262, 0.0465) | (0.0, 0.387) |

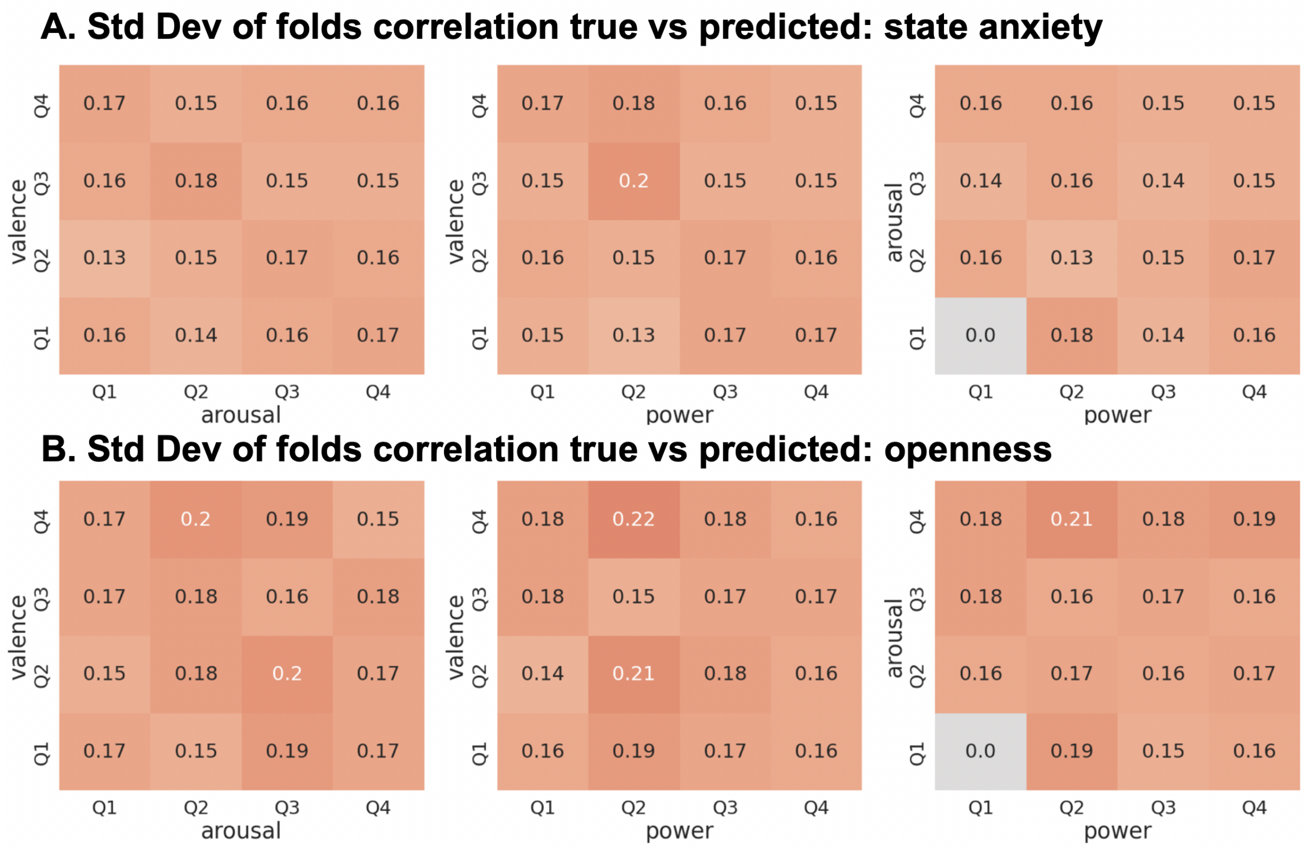


**Figure 7:** Standard deviation of correlation against consensus annotation of prediction (within all folds)

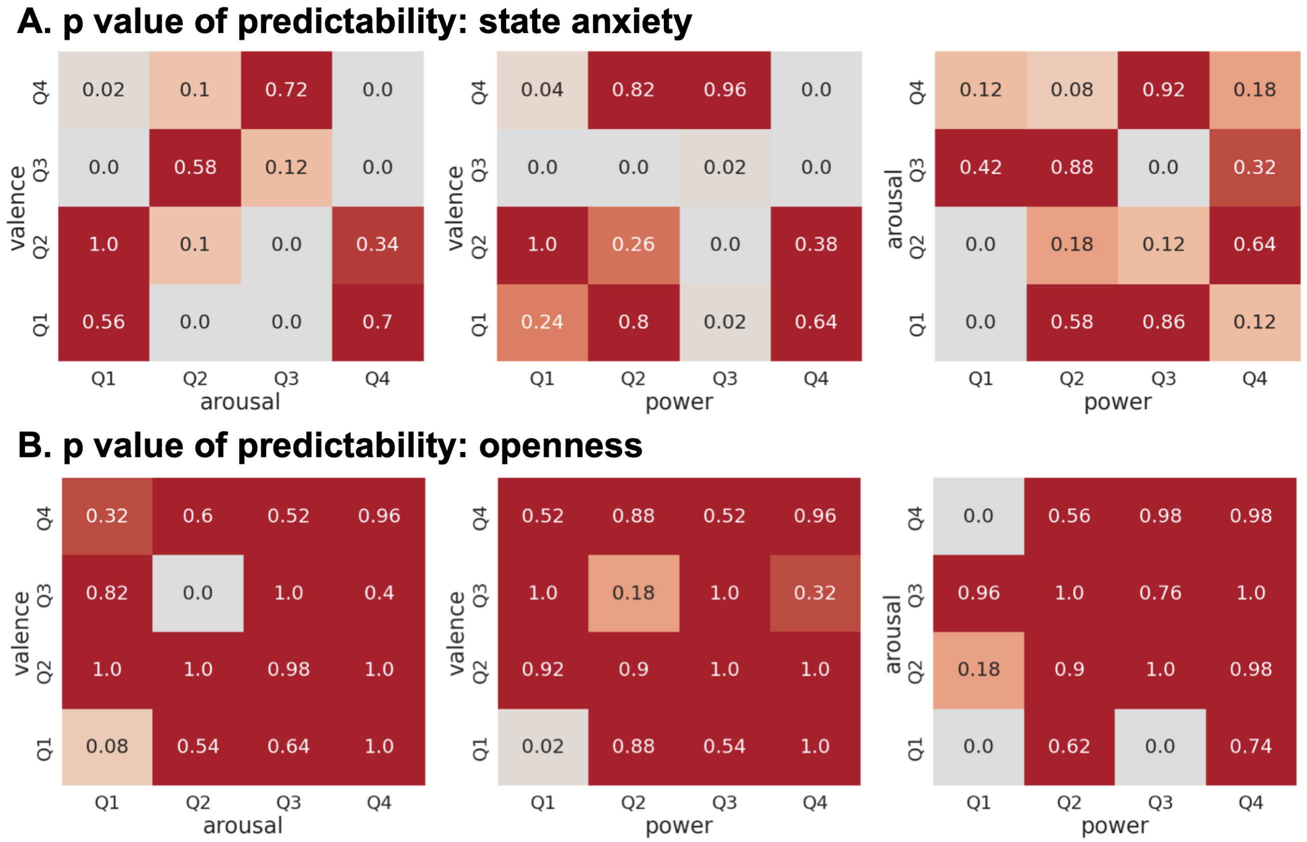


**Figure 8:** Prediction p-values using emotion-informed gradients

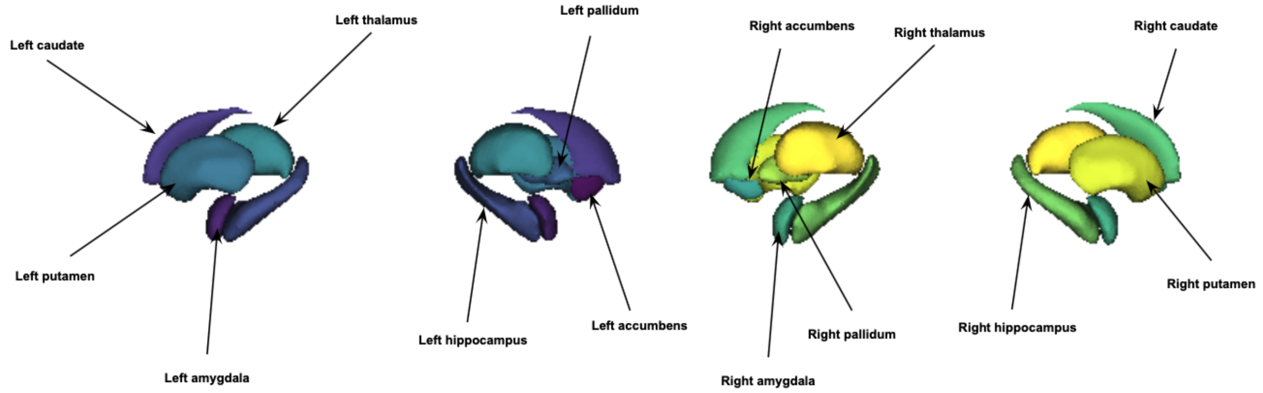


**Figure 9:** Legend for subcortical regions
